# The Crucial Role of CTCF in Mitotic Progression during Early Development of Sea Urchin

**DOI:** 10.1101/2023.05.26.542381

**Authors:** Kaichi Watanabe, Megumi Fujita, Kazuko Okamoto, Hajime Yoshioka, Miki Moriwaki, Hideki Tagashira, Akinori Awazu, Takashi Yamamoto, Naoaki Sakamoto

## Abstract

CCCTC-binding factor (CTCF), an insulator protein with 11 zinc fingers, is enriched at the boundaries of topologically associated domains (TADs) in eukaryotic genomes. In this study, we isolated and analyzed the cDNAs encoding HpCTCF, the CTCF homolog in the sea urchin *Hemicentrotus pulcherrimus*, to investigate its expression patterns and functions during early development of sea urchin. HpCTCF contains nine zinc fingers corresponding to fingers 2–10 of the vertebrate CTCF. The expression pattern analysis revealed that *HpCTCF* mRNA was detected at all developmental stages and in the entire embryo. Upon expressing the HpCTCF-GFP fusion protein in early embryos, we observed its uniform distribution within interphase nuclei. However, during mitosis, it disappeared from the chromosomes and subsequently reassembled on the chromosome during telophase. Moreover, the morpholino-mediated knockdown of HpCTCF resulted in mitotic arrest during the morula-to-blastula stage. Most of the arrested chromosomes were not phospholylated at serine 10 of histone H3, indicating that mitosis was arrested at the telophase by HpCTCF depletion. Furthermore, impaired sister chromatid segregation was observed using time-lapse imaging of HpCTCF-knockdown embryos. Thus, HpCTCF is essential for mitotic progression during the early development of sea urchins, especially during the telophase-to-interphase transition. However, the normal development of pluteus larvae in CRISPR-mediated HpCTCF knockout embryos suggests that disruption of zygotic HpCTCF expression has little effect on embryonic and larval development.

## Introduction

Eukaryotic genomes are organized into higher-order hierarchical chromatin structures, such as chromosome territories, A/B compartments, and topologically associated domains (TADs) (Cremer, & Cremer, 2001; Szabo et al., 2019; Dixon et al., 2012; Racko et al., 2019). The human genome contains approximately 2,000 TADs in a nucleus, with each TAD estimated to have an average size ranging from 500 kb to 1 Mb (Dixon et al., 2012). Genomic DNA sequences within the same TAD have a greater frequency of physical contact, enabling targeted communication between genes and nearby cis-regulatory elements such as enhancers for appropriate expression (Sexton et al., 2007; Dowen et al., 2014). This organization promotes precise gene expression.

CCCTC-binding factor (CTCF) was originally isolated as a transcription factor that specifically binds to the promoter-proximal regulatory region of c-Myc gene (Lobanenkov et al.,1990). CTCF is a DNA-binding protein with 11 zinc fingers that recognizes long and diverse DNA sequences (Kim et al., 2007; Ohlsson, Renkawitz, & Lobanenkov, 2001) and is involved in multiple functions, including transcriptional regulation, insulator function, and splicing control (Alharbi, Schmitz, Bailey, & Rasko, 2021; Braccioli & De Wit, 2019). Furthermore, CTCF has been reported to be enriched in TAD boundaries, and the constitution of TAD is regulated by CTCF and cohesin complexes (Merkenschlager & Nora, 2016; Parelho et al., 2008; Wendt et al., 2008). The CTCF is conserved in bilateral animals (Heger, Marin, Bartkuhn, Schierenberg, & Wiehe, 2012).

The requirement of CTCF for animal development has been investigated in several species (Fudenberg & Nora, 2021). Transgenic mice expressing RNAi targeting CTCF cannot develop into blastocyst stage embryos (Fedoriw et al., 2004). However, zygotic CTCF knockout mice can develop to the blastocyst stage but show lethality at the preimplantation stage (Moore et al., 2012). Furthermore, recent work has revealed that both maternal and maternal-zygotic CTCF knockout mice can develop up to the blastocyst stage. These findings indicate that maternal CTCF is not essential for early development (Andreu et al., 2022). In zebrafish, zygotic CTCF knockout embryos can develop normally through gastrulation and organogenesis until the pharyngeal stage. This is likely attributed to the presence of maternal CTCF. However, these embryos exhibit lethality at a later stage of development (Franke et al., 2022). Further, morpholino-mediated knockdown has revealed severe morphological defects resulting in lethality (Carmona-Aldana et al., 2018). In Drosophila, a meternal-zygotic knockout of dCTCF also showed reduced viability, depending on the genetic background (Kyrchanova et al., 2021).

Sea urchins are invertebrate deuterostomes that diverged during the early period of deuterostome evolution and have been used as model organisms for cell and developmental biology. During the cleavage stage of sea urchin development, embryos exhibit rapid cell division comprising primarily of the S and M phases. Subsequently, embryos undergo the formation of three germ layers, ectoderm, mesoderm, and endoderm, under the control of the gene regulatory network (Davidson et al., 2002; Oliveri & Davidson, 2004; Oliveri, Tu, & Davidson, 2008). The transparency of sea urchin embryos is beneficial for observing the intracellular behavior of molecules. As the microinjection technique is available for fertilized eggs, the expression of fluorescent proteins by mRNA microinjection and morpholino-mediated knockdown has been performed to analyze gene function. Furthermore, genomic sequences have been deciphered in a few sea urchin species and are available in public databases (Sea Urchin Genome Sequencing Consortium, 2006; Kinjo, Kiyomoto, Yamamoto, Ikeo, & Yaguchi, 2018; Davidson et al., 2020). Zinc-finger nuclease (ZFN), transcription activator-like effector nuclease (TALEN), and CRISPR-Cas9 are genome editing technologies that have recently emerged and revolutionized the analysis of gene function (Hosoi, Sakuma, Sakamoto, & Yamamoto, 2014; Lin & Su, 2016; Ochiai et al., 2010; Liu, Awazu, Sakuma, Yamamoto, & Sakamoto, 2019).

In this study, we analyzed the expression pattern and function of the sea urchin homolog of CTCF (HpCTCF) during early development of the sea urchin, *Hemicentrotus Plucherrimus*. Four *HpCTCF* cDNA variants encoding identical amino acid sequences were identified. *HpCTCF* mRNA was expressed at all developmental stages and throughout the embryonic region. Knockdown of HpCTCF by Morpholino antisense oligonucleotides resulted in the impairment of sister chromatid segregation and mitotic arrest of the cell cycle during telophase. However, the CRISPR-mediated knockout of HpCTCF did not result in any developmental anomalies. These results indicate that HpCTCF is essential for mitotic progression during early development of sea urchin.

## Materials and methods

### Embryo culture

Gametes of the sea urchin *H. pulcherrimus* were obtained by coelomic injection of 0.55 M KCl, and fertilized eggs were cultured in filtered sea water at 16°C.

### Cloning of *HpCTCF* cDNA

HpCTCF cDNA was originally isolated from the gastrula cDNA library constructed using the λgt11 vector. Total RNA was isolated from the mesenchyme blastula embryos using the acid guanidine phenol chloroform (AGPC) extraction method as described by Chomczynski and Sacchi (1987), and five micrograms of the total RNA were used for cDNA synthesis by SuperScript III reverse transcriptase (Invitrogen, USA). HpCTCF cDNA was amplified by RT-PCR with TaKaRa LA-taq (TaKaRa Bio Inc., Japan) using the following primers (5’-GTTCGCCATTCATTTGAGTGGATATTTC-3’ and 5’- CTGATCAGGTAAACTCTAAATCAGATTCC- 3’).

PCR products were electrophoresed on a 1% agarose gel, and approximately 5.0 kb of the PCR product was excised from the gel and subcloned into the pCR^®^4-TOPO^®^ vector using the TOPO TA Cloning^®^ Kit (Invitrogen, USA). The nucleotide sequences of the cDNA ends were determined by dideoxy chain termination chemistry using a Thermo Sequenase™ Cycle Sequencing Kit (USB Corporation, USA) and an LI-COR 4000L DNA sequencer (Aloka, Japan). The nucleotide sequence of the internal region was determined by BigDye^®^ Terminator v3.1 Cycle Sequencing Kit (Applied Biosystems, USA) and ABI PRISM^®^3100 Genetic Analyzer (Applied Biosystems, USA).

### Northern blot analysis

To prepare the antisense probe, part of the coding sequence (nucleotides 1069-2480) was excised by *Pst*I digestion from *HpCTCF* cDNA and subcloned into *Pst*I site of the pBluescript SK- vector. The antisense probe was labeled with digoxigenin-11-UTP (Roche, Germany) by MEGAscript^®^ T7 Kit (Ambion, USA). The labeling process involved utilizing a *Bam*HI-digested plasmid as a template, followed by purification through lithium chloride precipitation as described in the manufacturer’s instructions.

Total RNA was extracted from *H. pulcherrimus* embryos at various developmental stages using the AGPC extraction. Three micrograms of each RNA sample were electrophoresed on a denaturing 1% agarose gel containing formaldehyde and transferred to a Nytran^®^ N membrane (Whatman, USA). Hybridization was carried out in a CHURCH buffer (1% BSA, 1 mM EDTA, 0.5 M NaHPO_4_ (pH 7.2), 7% SDS) (Church & Gilbert, 1984) containing 10 ng/ml of the antisense RNA probe overnight at 65 °C. After washing the membrane with 1×SSC/1% SDS at 65 °C, hybridized probes were recognized using an anti-digoxigenin antibody conjugated to alkaline phosphatase (Roche, Germany) and detected by chemiluminescence produced by the dephosphorylation of CSPD (TROPIX, USA) using X-ray film (FUJIFILM, Japan).

### Whole mount in situ hybridization

Whole mount in situ hybridization was performed as described by Minokawa et al. (2004). To prepare the probe, a part of the coding sequence (nucleotides 1069-2480) was excised by *Pst*I digestion from *HpCTCF* cDNA and subcloned into *Pst*I site of the pBluescript SK- vector. The antisense probe was labeled with digoxigenin-11-UTP (Roche, Germany) by MEGAscript^®^ T7 Kit (Ambion, USA) using *Bam*HI digested plasmid as a template. To label the sense probe, the MEGAscript^®^ T3 Kit (Ambion, USA) and *Xho*I-digested plasmid were used.

### Expression of HpCTCF-GFP fusion protein

To express HpCTCF-GFP fusion protein in the *H. pulcherrimus* embryos, the HpCTCF coding sequence was subcloned into a pGreenLantern2-derived plasmid to express the HpCTCF-GFP fusion protein in *H. pulcherrimus* embryos. After linearization of the plasmid, 5’-capped mRNA encoding the HpCTCF-GFP fusion protein was synthesized using the mMESSAGE mMACHINE™ T7 Trinscription Kit (Ambion, USA) and purified by phenol/chloroform extraction and isopropanol precipitation, as described in the manufacturer’s instructions.

The *HpCTCF-GFP* mRNA was dissolved at 250 ng/µl in 40% glycerol and microinjected into the fertilized eggs of *H. pulcherrimus* as described by Rast (2000) with some modifications. Time-lapse analysis of GFP fluorescence was performed using the Olympus IX81 microscope and Metamorph software (Olympus, Japan).

### Knockdown by antisense morpholino oligonucleotide

Morpholino oligonucleotides complementary to the translation start site of HpCTCF (HpCTCF-MO1: 5’-AGGTTGGTCTGTGTTTTCATCCATG-3’ and HpCTCF-MO2: 5’-CATCCATGGTTCCCTCGTCTTATGA-3’) were synthesized by Gene Tools (Corvallis, USA), and the standard control morpholino oligonucleotide (5’- CCTCTTACCTCAGTTACAATTTATA-3’) was purchased from Gene Tools (Corvallis, USA). These morpholinos were dissolved in 40% glycerol, and 2 pl of morpholino solution was microinjected into the fertilized eggs of *H. pulcherrimus*.

### Immunohistochemistry

Sea urchin embryos were fixed in fixative III (4% paraformaldehyde, 32.5% filtered seawater, 32.5 mM MOPS (pH 7.0), 162.5 mM NaCl) at 4 °C for 16 h. After three washes with 1×PBS, the embryos were stored at -20 °C in ethanol. The fixed embryos were permeabilized with 0.5% Triton X-100 for 20 min at room temperature after three washes with 1×PBS. The embryos were subsequently blocked with 1×PBS with 1% BSA for an hour at room temperature, and then incubated with Anti-Histone H3 (phospho S10) antibody (mAbcam 14955) diluted at 1:2000 in 1×PBS with 1% BSA overnight at 4 °C. After three washes with 1×PBS, the embryos were incubated with Alexa Fluor 555- conjugated goat anti-rabbit polyclonal antibody (abcam150078) diluted at 1:2000 in 1×PBS with 1% BSA for 2 h at room temperature. After three washes with 1×PBS, the samples were mounted with SlowFade™ Gold Antifade Mountant with DAPI (Molecular Probes) and observed using an LSM700 confocal microscope (Zeiss, Germany).

### Knockout by CRISPR-Cas9

The CRISPR-Cas9-mediated knockout of *HpCTCF* was performed as described by Liu et al. (2019). The nucleotide sequence of the oligonucleotides used in the preparation of sgRNA#1 was 5’-GTAATACGACTCACTATAGGGAGGGGAGCCTGGTACTGGTTTTAGAGCTAGAAATAG-3’, that of sgRNA#2 was 5’-GTAATACGACTCACTATAGGTAGTGGGTGTGGTCATGGGTTTTAGAGCTAGAAATAG-3’, that of sgRNA#3 was 5’-GTAATACGACTCACTATAGGGGATGGCGCATCGGGGATGTTTTAGAGCTAGAAATAG-3’, and that of sgRNA#4 was 5’-GTAATACGACTCACTATAGGCATTAGGATTGCTGATGTGTTTTAGAGCTAGAAATAG-3’. Twenty-four hours after the microinjection of 750 ng/µl of Cas9 mRNA and 150 ng/µl of sgRNA into fertilized eggs, 20 embryos were collected, genomic DNA was isolated, and hetero-duplex mobility assay (HMA) was carried out using four primer sets (5’- TTTGGCAACATGTAACAGACTTGCCC-3’ and 5’-GTCTGCTCTTCACCAAGATCGTGGAC-3’ for sgRNA#1-target site; 5’- TTTGGCAACATGTAACAGACTTGCCC-3’ and 5’-TGAGACGGGTTGTAGCTCTGCATAGG -3’ for sgRNA#2-target site; 5’-CATCATGGACATGTCTGATAATTCTCTG-3’ and 5’-GAAGGCTCTGGTGTGTGGACTTGCC-3’ for sgRNA#3-target site; 5’- GTGGTAAAGCAGGAGATAGGTGAGG-3’ and 5’- CATATCTCACAAGATGCCTTAATCTC-3’ for sgRNA#4-target site). The PCR products were separated on a 0.5% agarose gel supplemented with the resolution enhancer Loupe 4K/20 (GelBio LLC, Japan) in 1×TAE buffer.

For sequencing analysis, the PCR products were subcloned into *Eco*RV site of pBluescript SK- vector using In-Fusion® HD Cloning Kit (TaKaRa Bio, Japan) and sequenced with the M13 forward primer (5’-GTAAAACGACGGCCAG-3’) using a BigDye Terminator v3.1 Cycle Sequencing Kit (Thermo Fisher Scientific).

## Results

### Nucleotide sequence of *HpCTCF*

We first isolated *HpCTCF* cDNA by screening the *H. pulcherrimus* gastrula cDNA library. Based on the nucleotide sequences of the isolated cDNA, two primers were designed in both 5’- and 3’-untranslated regions and performed RT-PCR using total RNA extracted from the mesenchyme blastula embryos. Four types of *HpCTCF* cDNAs were obtained by cloning approximately 5 kb of the PCR products (Fig. 1a), and the nucleotide sequences of these *HpCTCF* cDNA have been submitted to the DNA Data Bank of Japan (DDBJ) database under accession numbers LC767350, LC7673501, LC767352, and LC767353. The nucleotide sequences of these *HpCTCF* cDNA shared the identical nucleotide sequences in 5’-untranslated region (UTR) and the coding regions. However, the 3’-UTR sequences displayed distinct compositions, suggesting the presence of different exons owing to alternative splicing. Searching the *H. pulcherrimus* Genome and Transcriptome database (HpBase, http://cell-innovation.nig.ac.jp/Hpul/), revealed a single scaffold containing the *HpCTCF* cDNA sequence, which included exons corresponding to each part of 3’-UTR (Fig. 1b). Introduction of GFP mRNA fused with each type of 3’-UTR into sea urchin embryos did not result in any observable differences in GFP fluorescence distribution throughout the embryos (data not shown). This suggests that the 3’- UTRs are unlikely to contribute to the localization of *HpCTCF* mRNA.

**Figure 1.**
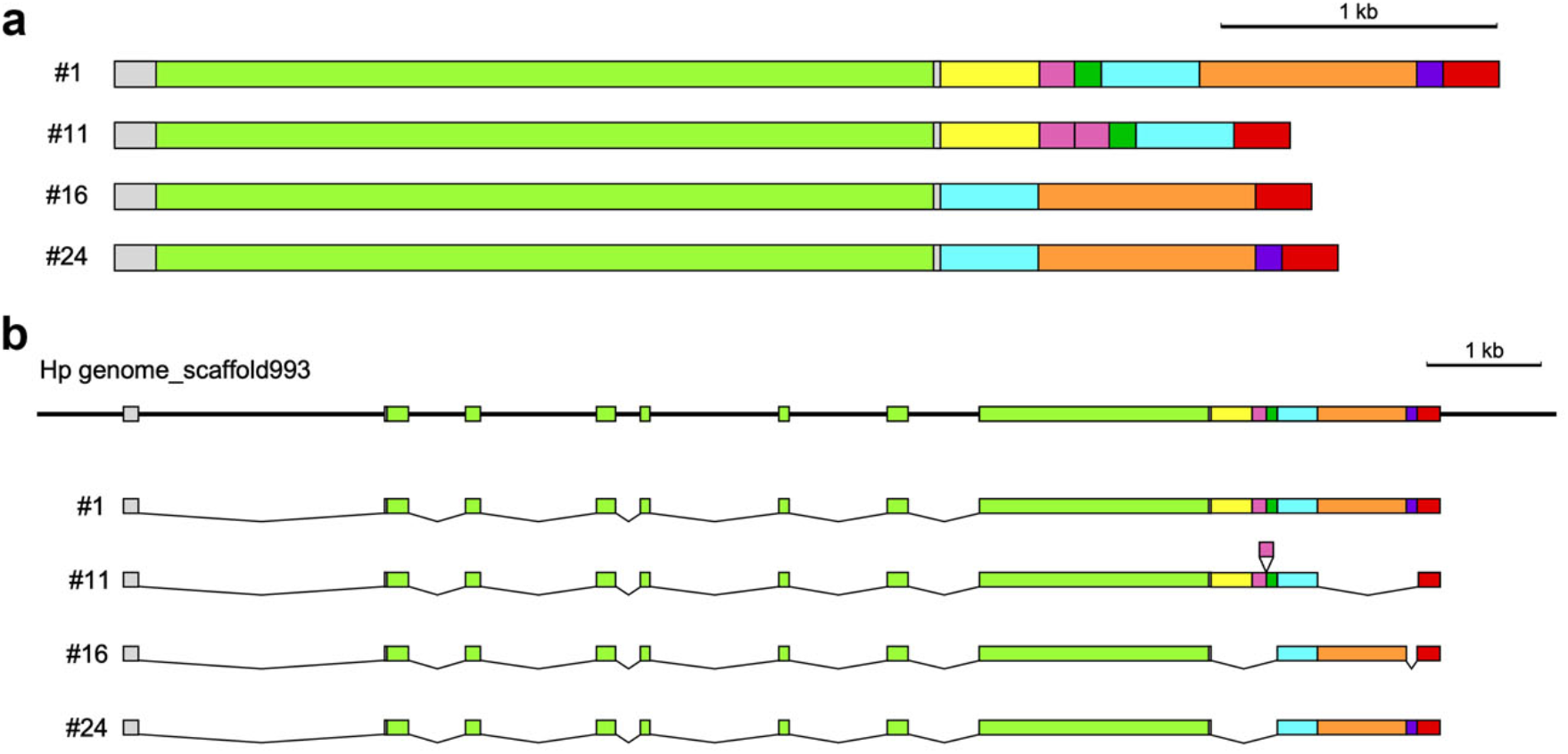
Schematic structures of HpCTCF cDNA (a) Four clones of *HpCTCF* cDNA isolated in this study are indicated. Coding regions are indicated by green boxes, and different nucleotides sequences are indicated by different colors. (b) Correspondence of each region of *HpCTCF* cDNA to the *HpCTCF* gene.

These cDNAs encoded an identical polypeptide 941 amino acids in length, with a single AT-hook motif and multiple C2H2-type zinc finger motifs (Fig. 2a). The amino acid sequence of HpCTCF exhibited less than 30% identity to human and Drosophila CTCF. However, the zinc finger region of HpCTCF was well conserved, with 51.3% and 47.8% identity, as well as 84.4% and 80.5% similarity, to the corresponding regions in human and Drosophila CTCF, respectively. Furthermore, although mammalian and Drosophila CTCF contain 11 zinc finger motifs, only nine are present in HpCTCF (Fig. 2a). Interspecific comparison of zinc finger motifs revealed that the zinc finger motifs of HpCTCF (ZF1 to ZF9) are homologous to ZF2 to ZF10 in vertebrate and Drosophila CTCF (Fig. 2b). Moreover, a homology search using the Lipman–Pearson method revealed that of the nine zinc finger motifs of HpCTCF, ZF5 to ZF7 showed higher homology (optimized score) to protostome animals’ (Drosophila and nematode) CTCF than human CTCF. This suggests that HpCTCF may possess an ancestral type of structure and function.

**Figure 2.**
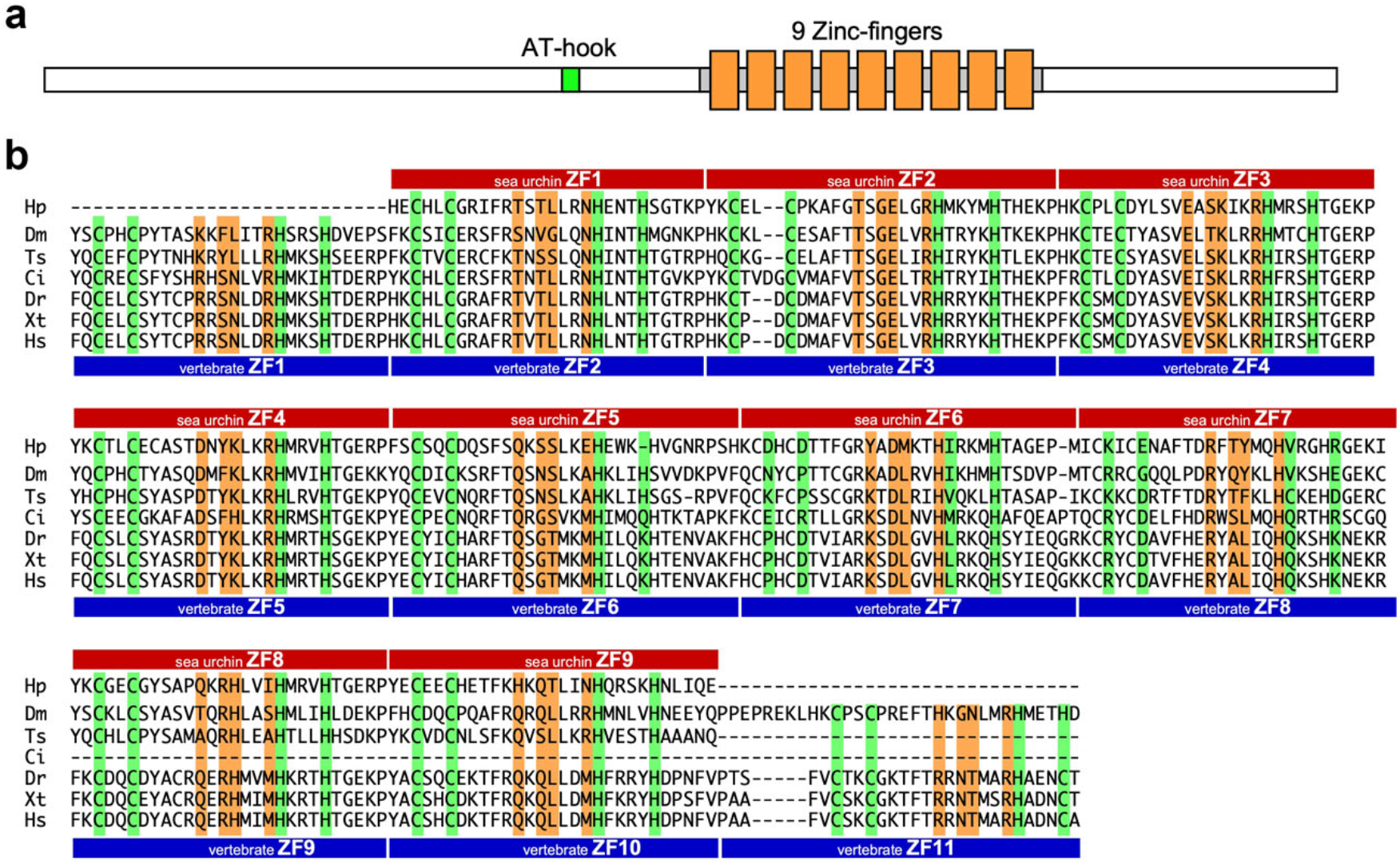
Structure and amino acid sequence of HpCTCF. (a) Schematic illustration of HpCTCF proein. Zinc finger motifs are indicated by orange boxes, and AT-hook motif is indicated by green box. (b) Alignment of aminoacid sequences of zinc finger domains from various oganisms. Red and blue boxes indicate the positions of zinc finger motifs in sea urchin CTCF (HpCTCF) and vertebrate CTCF, respectively. Hp, zinc finger domain of HpCTCF; Dm, that of *Drosophila melanogaster* CTCF (NP_648109); Ts, that of *Trichinella spiralis* (basal nematode) CTCF (KRY34076); Ci, that of *Ciona intestinalis* (tunicate) CTCF (NP_001104593), Dr, that of *Danio rerio* (zebrafish) CTCF (NP_001001844); Xt, that of Xenopus tropicalis CTCF (`NP_001116268`); Hs, that of *Homo sapiens* (human) CTCF (NP_006556). Cysteins (C) and histidines (H) involved in the zinc finger formation are highlighted by green color, and residues involved in DNA recognition are highlighted by orange color.

### Expression patterns of *HpCTCF* mRNA

To analyze the temporal expression patterns of *HpCTCF* mRNA, we performed northern blot analysis using total RNAs isolated from *H. pulcherrimus* embryos at various developmental stages (Fig. 3). Maternal *HpCTCF* mRNA was present in unfertilized eggs. The amount of *HpCTCF* mRNA was maintained, with a slight reduction during the cleavage stage, and then increased and reached a maximum level at the hatched blastula stage. High levels of *HpCTCF* mRNA expression were maintained during the blastula stage, but the amount of *HpCTCF* mRNA decreased from the gastrula to the pluteus stage.

**Figure 3.**
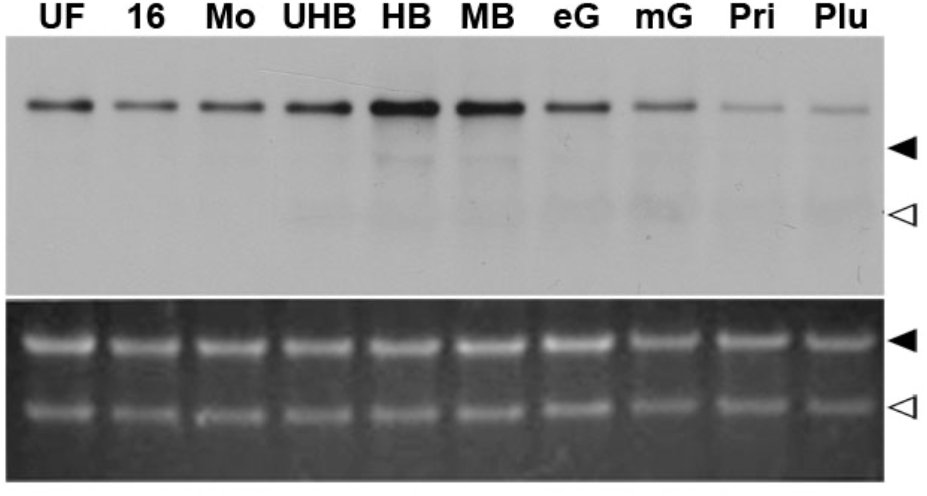
Northern blot analysis of *HpCTCF* mRNA. Five micrograms of total RNA extracted from the embryos at various developmental stages were electrophoresed. The upper panel shows northern blot analysis using an antisense probe complementary to *HpCTCF* mRNA, and the lower panel shows total RNA stained with ethidium bromide. Closed triangles indicate the position of 26S rRNA (4.2 kb) and open triangles indicate 18S rRNA (2 kb). UF, unfertilized egg; 16, 16-cell stage; Mo, morula; UHB, unhatched blastula; HB, hatched blastula; MB, mesenchyme blastula; eG, early gastrula; mG, mid-gastrula; Pri, prism larvae; Plu, pluteus larvae.

The spatial expression patterns of *HpCTCF* mRNA were analyzed using whole mount in situ hybridization (Fig. 4). Corresponding to the results of northern blot analysis, *HpCTCF* mRNA was detected throughout all developmental stages. Furthermore, *HpCTCF* mRNA was detected in the entire embryonic region at all developmental stages, suggesting a ubiquitous role of HpCTCF in sea urchin development.

**Figure 4.**
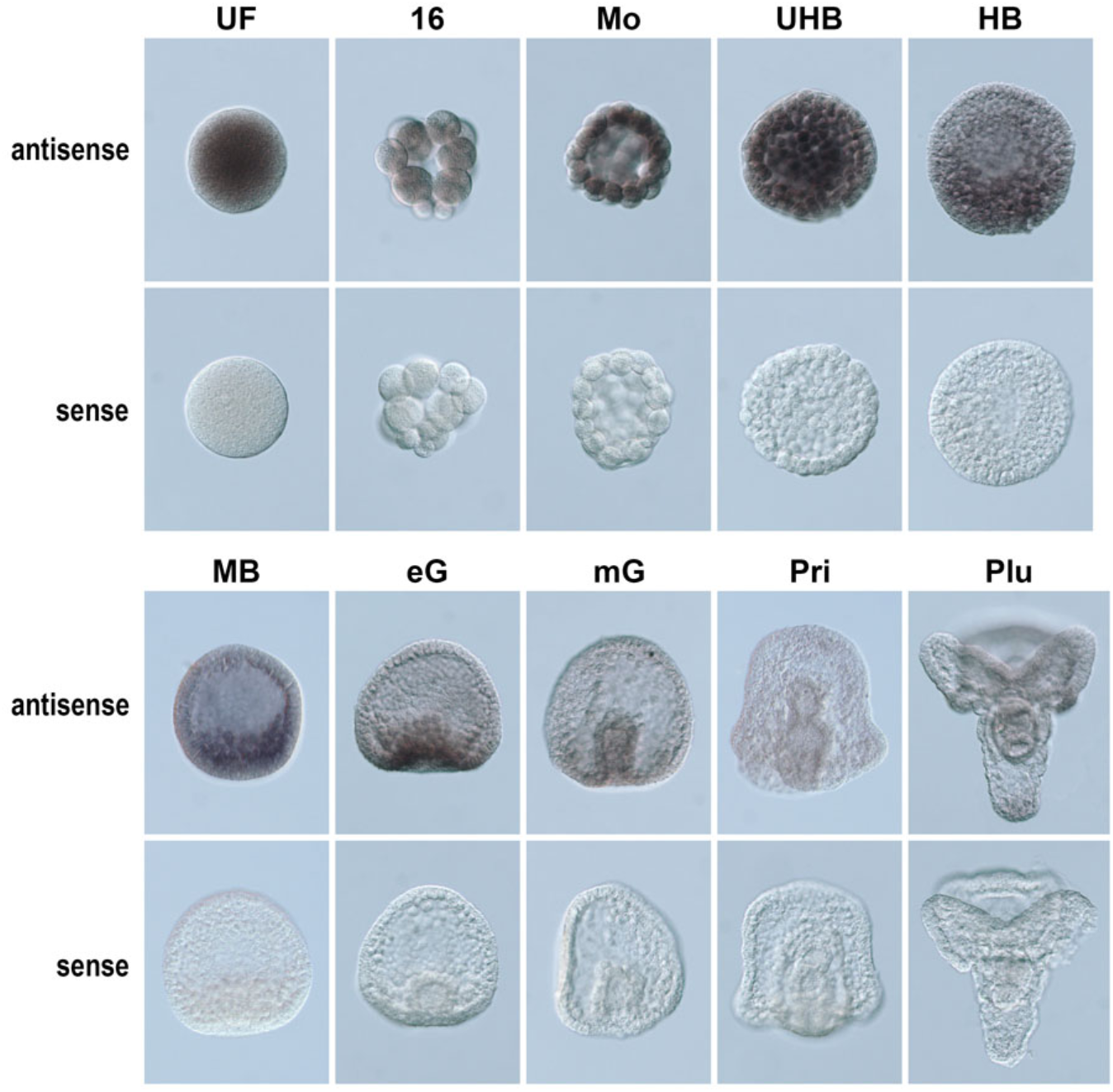
Whole mount in situ hybridization analysis of *HpCTCF* mRNA. Whole-mount in situ hybridization was performed using an antisense probe complementary to *HpCTCF* mRNA or a sense probe with a sequence identical to that of *HpCTCF* mRNA. UF, unfertilized egg; 16, 16-cell stage; Mo, morula; UHB, unhatched blastula; HB, hatched blastula; MB, mesenchyme blastula; eG, early gastrula; mG, mid- gastrula; Pri, prism larvae; Plu, pluteus larvae.

### Subcellular localization of *HpCTCF*

To examine the role of HpCTCF during development, we analyzed its subcellular localization of HpCTCF protein. The mRNA encoding the HpCTCF-GFP fusion protein was microinjected into fertilized eggs of *H. pulcherrimus* and GFP fluorescence was analyzed through time-lapse imaging during the cleavage stage (Fig. 5). HpCTCF-GFP was distributed throughout the nucleus, and this nuclear fluorescence disappeared approximately 15 min before cell division but was restored after cell division (Fig. 5a).

**Figure 5.**
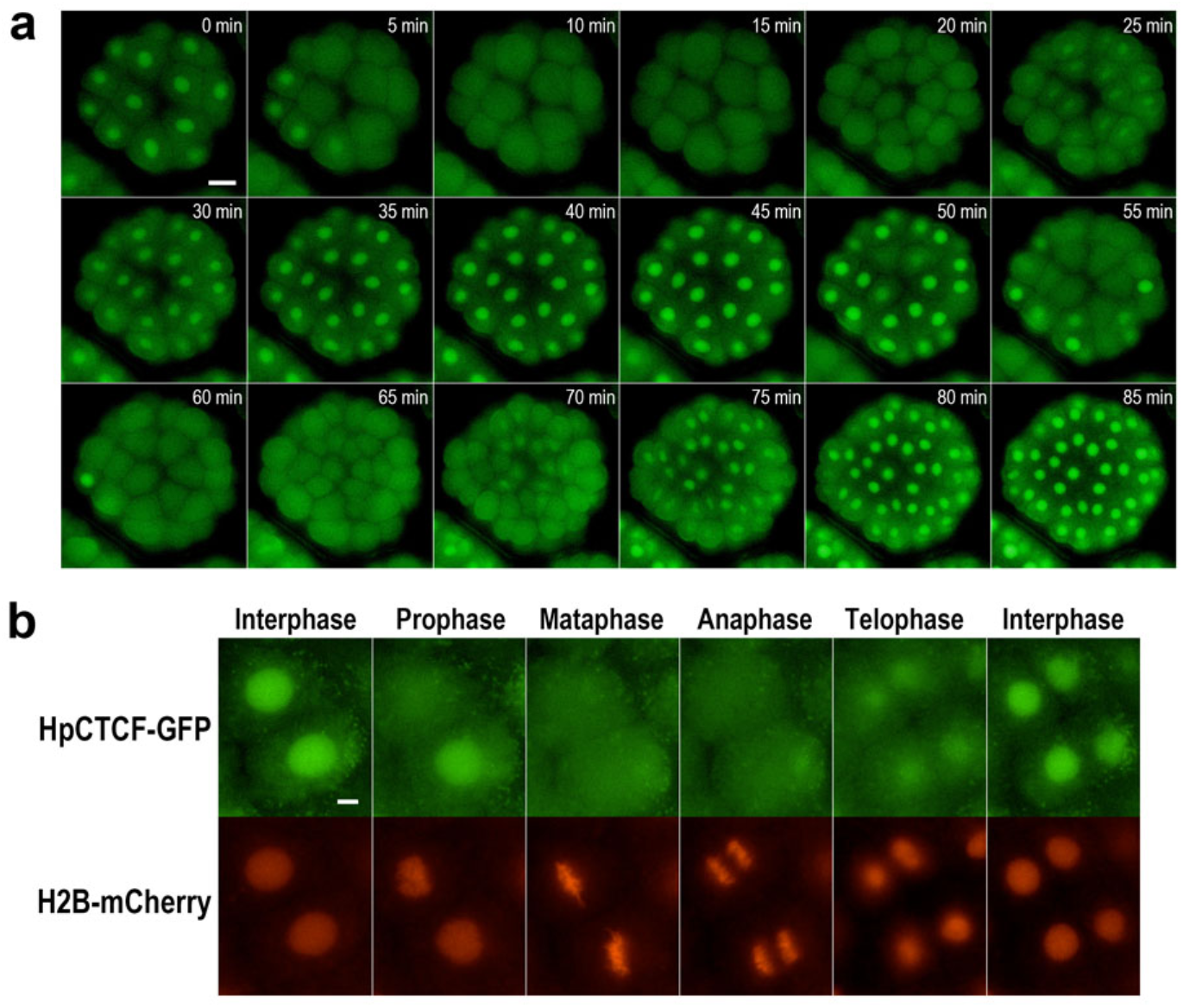
Cell cycle-dependent behavior of HpCTCF. (a) Time-lapse analysis of the behavior of HpCTCF-GFP fusion protein in the entire embryo. Scale bar: 20 µm. (b) Time-lapse analysis of HpCTCF-GFP and H2B-mCherry during the cell cycle of the morula embryo. Scale bar: 5 µm.

For a detailed analysis of the subcellular behavior of HpCTCF, *H2B-mCherry* mRNA was coinjected with *HpCTCF-GFP* mRNA and analyzed using time-lapse imaging. As shown in Fig. 5b, HpCTCF-GFP was detected in the interphase nuclei; however, HpCTCF-GFP fluorescence started to decrease at the onset of chromosome condensation (prophase) and was not detected on chromosomes from metaphase to anaphase. HpCTCF-GFP fluorescence resumed at telophase, during which interphase nuclei were reconstructed. These results suggest that HpCTCF may play a role in the reorganization of interphase nuclei during sea urchin cell cycle development.

### Effects of HpCTCF knockdown on the early development

To analyze the functional role of HpCTCF in sea urchin development, a morpholino antisense oligonucleotide complementary to *HpCTCF* mRNA (HpCTCF-MO) was employed to inhibit the translation of HpCTCF. This was achieved by microinjecting HpCTCF-MO into fertilized eggs of *H. pulcherrimus*. Two HpCTCF-MOs (MO1 and MO2) were designed to target the translational initiation of both zygotically expressed *HpCTCF* mRNA and maternal *HpCTCF* mRNA.

To analyze the phenotype, the morphology of HpCTCF-MO-injected embryos was compared with that of embryos injected with a control morpholino. At the mesenchyme blastula stage, approximately 95% of control embryos exhibited normal morphology, characterized by distinct blastocoels and well defined primary mesenchyme cells (Fig. 6a and c). In contrast, approximately 90% of MO1-injected embryos and half of the MO2- injected embryos exhibited an abnormal appearance, characterized by an opaque blastocoel filled with disorganized cells (Fig. 6b and c). These abnormal embryos subsequently ceased to develop. The HpCTCF-MO phenotype was observed in a dose-dependent manner (Fig. 6d).

**Figure 6.**
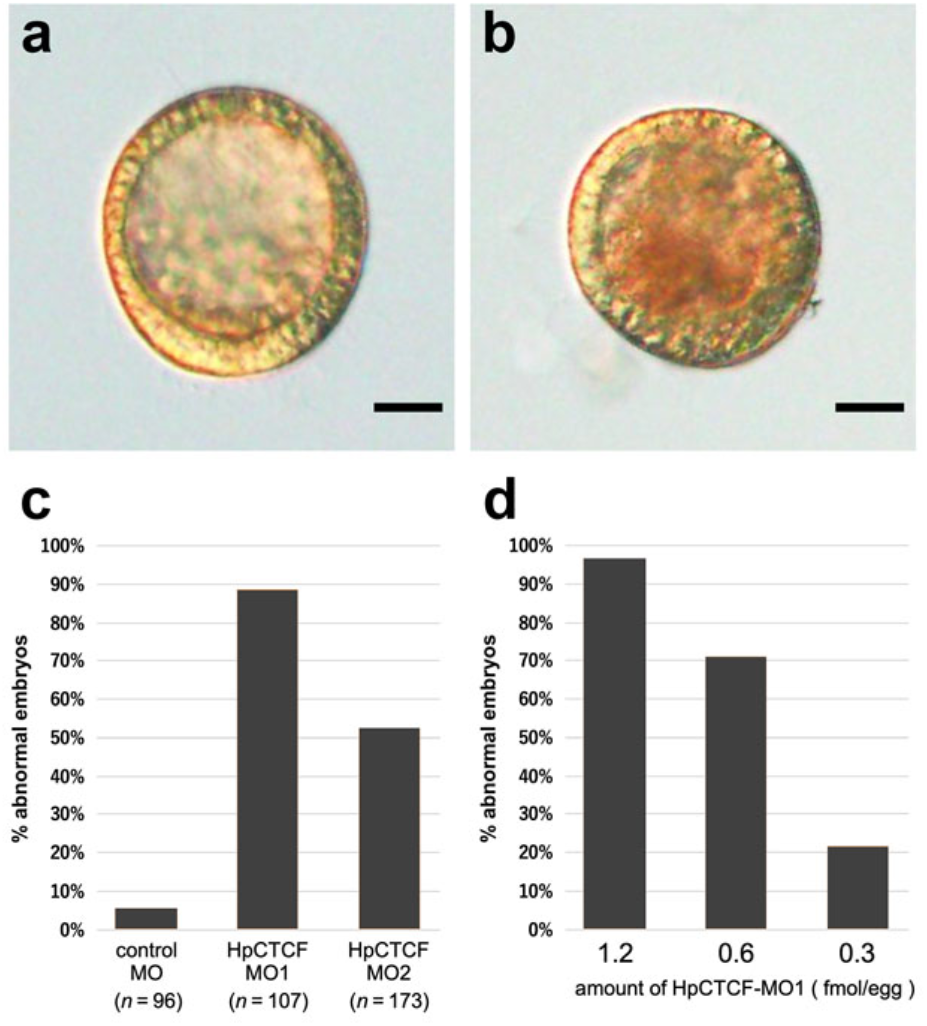
Effects of HpCTCF knockdown on the early development of sea urchin. (a) Embryos injected with control MO at 2.0 fmol/egg. (b) Embryos injected with HpCTCF-MO1 at 2.0 fmol/egg. Scale bar: 30 µm. (c) Proportion of abnormal embryos in control and HpCTCT-MO injected embryos. Each 2 fmol of MO was injected into each egg. (d) Dose-dependent effect of HpCTCF-MO1. HpCTCF-MO1 was injected at cocentrations indicated below the graph and the proportion of the abnormal embryos were scored.

To analyze this phenotype in detail, the chromosomal DNA of HpCTCF-MO1-injected embryos was stained with DAPI. Although the majority of cells in the control embryos exhibited normal interphase nuclei (Fig. 7a, left panel), several cells in the HpCTCF- MO1-injected embryos exhibited condensed chromosomal DNA, which may be due to mitotic arrest of their cell cycle (Fig. 7a, right panel).

**Figure 7.**
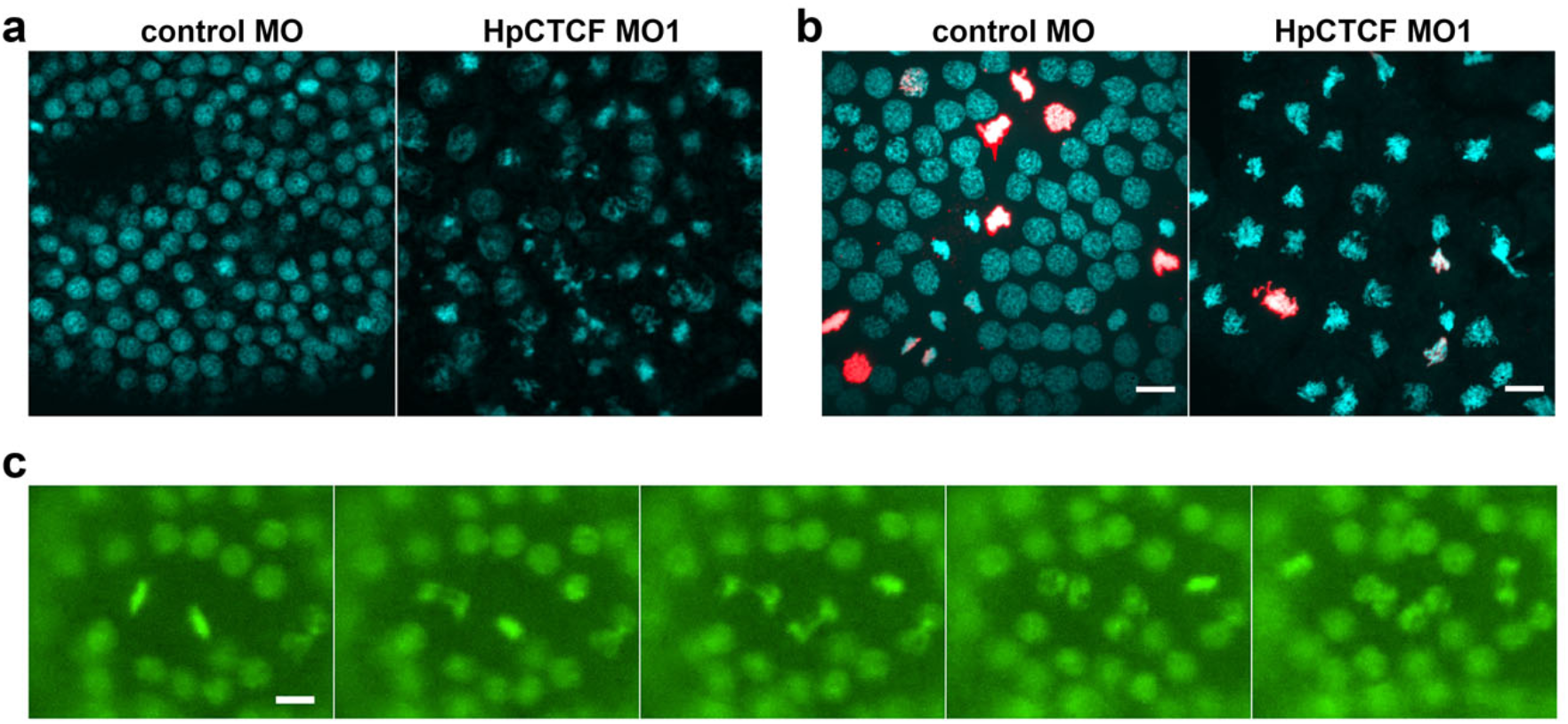
Chromosomes observed in the HpCTCF knockdown embryos. (a) DAPI staining of chromosomes in control MO and HpCTCF-MO1-injected embryos. (b) Immunohistochemistry of H3S10 phosphorylation in control MO and HpCTCF-MO1- injected embryos. (c) Time-lapse analysis of H2B-GFP in the HpCTCF-MO1-injected embryos.

The mitotic arrest caused by HpCTCF-MO was further analyzed by staining chromosomal DNA of HpCTCF-MO1-injected embryos with an anti-phospho-histone H3 (Ser10) antibody, which is a well-known mitosis marker. In control embryos, high-level deposition of H3S10 phosphorylation was detected on the entire mitotic chromosome from prophase to metaphase and at the tip of chromatids at anaphase, but not during telophase (Fig. 7b, left panel). In contrast, in HpCTCF-MO1-injected embryos, most of the condensed chromosomal DNA was not recognized by this antibody (Fig. 7b, right panel), suggesting that the condensed DNA resulting from HpCTCF knockdown was in telophase. Furthermore, when HpCTCF-MO1 was co-injected with *H2B-GFP* mRNA to visualize the chromosomal behavior of HpCTCF-MO1-injected embryos, we observed an impairment of sister chromatid segregation during the morula-to-blastula stages (Fig. 7c).

### Effects of CRISPR-Cas9-mediated knockout of HpCTCF

To elucidate the function of zygotically expressed HpCTCF, we performed a CRISPR- Cas9-mediated knockout of *HpCTCF*. We designed two sgRNAs, #1 and #2, which targeted the region encoding the N-terminal side of AT-hook motif. These sgRNAs were microinjected with *SpCas9* mRNA into fertilized eggs, genomic DNA was extracted at 24 h post fertilization (hpf) from 20 embryos, and mutations introduced by CRISPR-Cas9 were examined by heteroduplex mobility assay (HMA). A band shift was detected in embryos co-injected with *SpCas9* mRNA and either sgRNA, whereas embryos injected with *SpCas9* mRNA alone did not exhibit any band shift (Fig. 8a).

**Figure 8.**
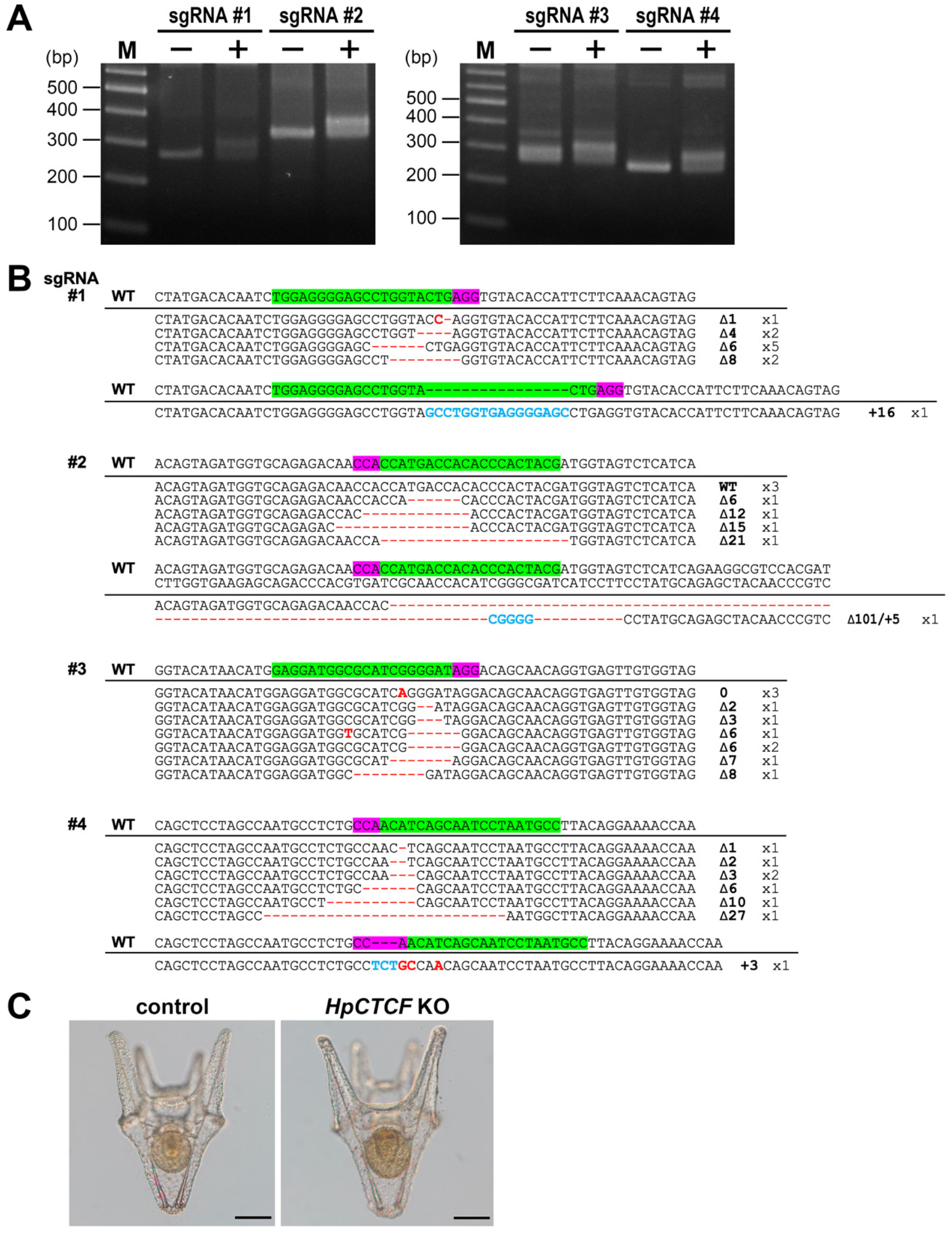
Effects of CRISPR-Cas9-mediated HpCTCF knockout on the early development of sea urchin. (a) Genotyping of sgRNA-injected embryos by HMA. Genomic DNA was extracted from 20 embryos injected with SpCas9 alone (-) or SpCas9/sgRNA (+) at 24 hpf, and the target sites ware analyzed by HMA. M: 100 bp ladder marker. (b) Sequence analysis of mutations induced by HpCTCF sgRNA#1, #2, #3 and #4. The wild-type sequences are shown on the top, with the PAM highlighted in magenta and the protospacer highlighted in green. Deletions, substitution, and insertion are indicated by red dashes, red letters, and blue letters, respectively. (c) Morphologies of control and HpCTCF sgRNA#1-injected knockout embryos. Scale bar: 100 µm.

PCR amplicons from HpCTCF-knockout embryos were subcloned and sequenced to analyze the type and efficiency of the induced mutations. Among the 11 sequenced clones from the sgRNA#1-injected knockout embryos, deletions (10 clones) and insertions (one clone) were observed, indicating that the mutation rate was 100%. However, five clones showed a 6-bp deletion, and the frameshift rate was 54.5% (Fig. 8b). In the sequencing analysis of sgRNA#2-injected knockout embryos, the mutation rate was 57.1%; however, all mutated clones had deletions in multiples of three nucleotides (Fig. 8b).

To maximize the CRISPR-mediated mutagenesis, we designed two additional sgRNAs, #3 and #4, targeting more N-terminal side of the HpCTCF coding sequence. These two sgRNA-injected embryos also exhibited effective HMA mutagenesis (Fig. 8a). Although sequencing analysis revealed that sgRNA#3-injected embryos showed a 100% mutation rate, all substitutions were silent mutations and the frameshift rate was 30%. Furthermore, sgRNA#4-injected embryos showed 100% mutation rate, but the frameshift rate was 37.5%.

All HpCTCF-knockout embryos developed normally until at least 10 days post-fertilization, and there was no considerable difference in size, morphology, or lethality between the control and knockout larvae (Fig. 8c). This result suggests that disturbances in zygotic HpCTCF expression have little effect on embryonic and larval development.

## Discussion

From an evolutionary perspective, sea urchins are an interesting animal model for biological research. In this study, we isolated cDNA encoding the CTCF homolog of the sea urchin *Hemicentrotus pulcherrimus,* HpCTCF. Although CTCF contains 11 zinc finger motifs in general, we found that HpCTCF contains only nine fingers that correspond to fingers 2–10 of vertebrate CTCF. The CTCF of the basal nematode *Trichinella spiralis* has 10 fingers corresponding to ZF1-10 of the vertebrate CTCF (Heger et al., 2009). The tunicate CTCF has eight fingers corresponding to ZF1-8 of the vertebrate CTCF. Notably, TAD has not been identified in tunicates (Satou et al., 2019). *C. elegans* has lost CTCF during evolution (Heger et al., 2009), and there is no strong TAD on its autosomes; however, dosage-compensated X chromosomes have TAD mediated by condensin DC (Crane et al., 2015; Kim et al., 2022). The existence of TAD in sea urchins has not been previously reported, suggesting that CTCF may change the number of zinc fingers depending on their role in TAD formation.

Comparing the amino acid sequence of HpCTCF with those of various animals revealed that ZF5-7 of HpCTCT exhibited greater homology to zinc fingers of protostome animal CTCF compared to those of human CTCF. Furthermore, ZF2 showed a higher optimal score in the homology search for Drosophila CTCF. This suggests that HpCTCF possesses an ancestral structure and ZF2 and ZF5-7 may be responsible for the basic role of HpCTCF. In humans, mutations in tumor-associated zinc-finger mutations have been reported in ZF3 (corresponding to HpZF2) and ZF7 (corresponding to HpZF6) (Filippova et al., 2002), suggesting that the basic role of CTCF may be associated with cell proliferation.

HpCTCF was ubiquitously expressed during the early development of sea urchins (Fig. 3 and 4), suggesting a ubiquitous role for HpCTCF during development. During the rapid cell cycle cleavage, HpCTCF exists uniformly within the interphase nuclei, but is undetectable on mitotic chromosomes, especially from prophase to anaphase (Fig. 5). The disappearance of CTCF from prometaphase chromosomes has been reported in HeLa cells (Oomen, Hansen, Liu, Darzacq, & Dekker, 2019), and the exclusion of CTCF from mitotic chromatin has been reported in WI-38 cells (Agarwal, Reisser, Wortmann, C., & Gebhardt, 2017). Furthermore, structures within the interphase nuclei, such as the A/B compartments and TAD, are lost during prometaphase (Naumova et al., 2013), and transcription levels decrease during mitosis (Palozola et al., 2017). Although the presence of TAD in the interphase nuclei of sea urchins remains uncertain, HpCTCF is believed to play an important role in these nuclei during the early development of sea urchins.

In the HpCTCF knockdown experiment, phosphorylation of serine 10 of histone H3 (H3S10) was analyzed by immunohistochemistry. H3S10 phosphorylation was detected in all mitotic chromosomes from prophase to metaphase and at the tip of the chromatids at anaphase, but not during telophase (Fig. 7b). H3S10 phosphorylation in sea urchins has been shown to follow the same pattern as in MCF-7 cells (Yan et al., 2016), indicating that H3S10 phosphorylation is conserved in sea urchins. HpCTCF knockdown resulted in mitotic arrest of the cell cycle, and most of the arrested condensed chromosomes in the knockdown embryos did not contain H3S10 phosphorylation, suggesting that the arrested mitotic chromosomes were in telophase. This may be due to the failure of post-mitotic accumulation of HpCTCF during telophase and the reorganization of interphase nuclei.

Furthermore, knockdown led to the impairment of sister chromatid segregation during anaphase. Although HpCTCF could not be detected during anaphase, the presence of CTCF on mitotic chromosomes, centrosomes, and midbodies has been previously reported in mammalian cells (Burke et al., 2005). Therefore, a small amount of HpCTCF may exist on mitotic chromosomes and spindles, and contribute to sister chromatid segregation. A recent report showed that the CRISPR-mediated knockdown of CTCF leads to disorganized tri/tetrapolar spindle formation and impairment of anaphase segregation in mouse B16-F1 cells (Chiu et al., 2023).

In the CRISPR-mediated knockout of HpCTCF, embryos developed into pluteus larvae with normal morphology (Fig. 8c). At pluteus larvae, it was reported that Br-dU positive proliferating cells are concentrated in the circumoral ectoderm including larval arms (Katow, Katow, Yoshida & Kiyomoto, 2021). However, because the amount of HpCTCF was remarkably reduced in pluteus larvae (Figs. 3 and 4), cell proliferation in pluteus larvae may not require HpCTCF. Alternatively, as the frameshift rate in the sgRNA-injected HpCTCF knockout was 54.5% or less, this efficiency may be insufficient to disrupt the contribution of HpCTCF to cell proliferation. Furthermore, as almost all sgRNA-injected embryos were viable in all CRISPR-mediated knockout experiments, it did not appear that only embryos without this phenotype survived.

TAD plays an important role in promoter-enhancer interactions. During animal development, TAD is formed during activation of the zygotic genome (Du et al., 2017; Hug, Grimaldi, Kruse, & Vaquerizas, 2017; Ke et al., 2017; Wike et al., 2021). In the sea urchin *S. purpuratus*, the initiation time of zygotic gene expression is 5–6 hours postfertilization (Gildor & Ben-Tabou de-Leon, 2015; Materna, Nam, & Davidson, 2010). Correspondingly, the amount of *HpCTCF* mRNA began to increase (Fig. 3), suggesting that HpCTCF plays a role in zygotic gene expression. However, CRISPR-mediated HpCTCF-knockout embryos developed normally in pluteus larvae (Fig. 8c), suggesting that HpCTCF in interphase nuclei may not be essential for gene expression. Arthropods, including Drosophila, have acquired several other insulator proteins (Heger, George, & Wiehe, 2013), and it has been reported that arthropods have TAD regulatory mechanisms mediated by factors other than CTCF, and that CTCF contributes to loops in limited gene areas (Van Bortle et al., 2014; Kaushal et al., 2021; Cavalheiro et al., 2023). As insulator proteins other than CTCF have also been reported in sea urchins (Cavalieri, Melfi, & Spinelli, 2013; Yajima, Fairbrother, & Wessel, 2012), the phenotype of CRISPR-mediated HpCTCF-knockout embryos may be normal.

Collectively, the sea urchin CTCF has an ancestral function and is involved in cell cycle progression during early development. This hypothesis is also supported by the analysis of single-cell RNA-sequencing data from embryos of *Strongylocentrotus purpuratus*, a closely related species of *H. pulcherrimus*, at developmental stages in which both dividing and non-dividing cells are present (Foster et al., 2020), where a significant positive correlation was observed between the expression of CTCF and the expression of cell cycle- and M-phase chromosome formation-related genes (Table S1). The role of sea urchin CTCF in the organization of interphase nuclei and transcriptional regulation remains unclear. Future studies using chromosome conformation capture technologies can uncover the detailed function of sea urchin CTCF in interphase nuclei.

## Author Contributions

NS conceived and designed the experiments. KW, MF, KO, HY, MM, HT and NS conducted the experiments. AA provided instructions. KW, AA and NS wrote the manuscript with support from all authors. TY supervised the work.

## Acknowledgements

We thank Dr. Masato Kiyomoto (Tateyama Marine Laboratory, Ochanomizu University) for supplying live sea urchins and sea water. This work was supported by Grant-in-Aid for Scientific Research (C) (JSPS KAKENHI Grant Number JP20K06602) to NS, Grants-in-Aid for Scientific Research (C) (JSPS KAKENH Grant Number JP21K06124) to AA, and JST, the establishment of university fellowships towards the creation of science technology innovation, Grant Number JPMJFS2129.

## Notes

### Competing Interest Statement

The authors have declared no competing interest.

## References

1. Agarwal, H., Reisser, M., Wortmann, C., Gebhardt, J. C. M. (2017). Direct observation of cell-cycle-dependent interactions between CTCF and chromatin. Biophysical Journal, 112, 2051–2055. doi: 10.1016/j.bpj.2017.04.018

2. Alharbi, A. B., Schmitz, U., Bailey, C. G., & Rasko, J. E. J. (2021). CTCF as a regulator of alternative splicing: New tricks for an old player. Nucleic Acids Research, 49, 7825–7838. doi: 10.1093/nar/gkab520

3. Andreu, M. J., Alvarez-Franco, A., Portela, M., Gimenez-Llorente, D., Cuadrado, A., Badia-Careaga, C., … Manzanares, M. (2022). Establishment of 3D chromatin structure after fertilization and the metabolic switch at the morula-to-blastocyst transition require CTCF. Cell Reports, 41, 111501. doi: 10.1016/j.celrep.2022.111501.

4. Braccioli, L., & De Wit, E. (2019). CTCF: A Swiss-army knife for genome organization and transcription regulation. Essays in Biochemistry, 63, 157–165. doi: 10.1042/EBC20180069

5. Burke, L. J., Zhang, R., Bartkuhn, M., Tiwari, V. K., Tavoosidana, G., Kurukuti, S., … Renkawitz, R. (2005). CTCF binding and higher order chromatin structure of the H19 locus are maintained in mitotic chromatin. EMBO Journal, 24, 3291–3300. doi: 10.1038/sj.emboj.7600793

6. Carmona-Aldana, F., Zampedri, C., Suaste-Olmos, F., Murillo-de-Ozores, A., Guerrero, G., Arzate-Mejía, R., … Recillas-Targa, F. (2018). CTCF knockout reveals an essential role for this protein during the zebrafish development. Mechanism of Development, 154, 51–59. doi: 10.1016/j.mod.2018.04.006

7. Cavalheiro, G. R., Girardot, C., Viales, R. R., Pollex, T., Cao, T. B. N., Lacour, P., … Furlong, E. E. M. (2023). CTCF, BEAF-32, and CP190 are not required for the establishment of TADs in early *Drosophila* embryos but have locus-specific roles. Science Advances, 9, eade1085. doi: 10.1126/sciadv.ade1085.

8. Cavalieri, V., Melfi, R., & Spinelli, G. (2013). The Compass-like locus, exclusive to the Ambulacrarians, encodes a chromatin insulator binding protein in the sea urchin embryo. PLoS Genetics, 9, e1003847. doi: 10.1371/journal.pgen.1003847

9. Chiu, K., Berrada, Y., Eskndir, N., Song, D., Fong, C., Naughton, S., … Stephens, A. D. (2023). CTCF is essential for proper mitotic spindle structure and anaphase segregation. bioRxiv, doi: 10.1101/2023.01.09.523293, 10 Jan 2023, preprint: not peer reviewed.

10. Chomczynski, P., & Sacchi, N. (1987). Single-step method of RNA isolation by acid guanidinium thiocyanate-phenol-chloroform extraction. Analytical Biochemistry, 162, 156–159. doi: 10.1006/abio.1987.9999

11. Church, G. M., & Gilbert, W. (1984). Genomic sequencing. Proceedings of the National Academy of Sciences of the United States of America, 81, 1991–1995. doi: 10.1073/pnas.81.7.1991

12. Crane, E., Bian, Q., McCord, R. P., Lajoie, B. R., Wheeler, B. S., Ralston, E. J., … Meyer, B. J. (2015). Condensin-driven remodelling of X chromosome topology during dosage compensation. Nature, 523, 240–244. doi: 10.1038/nature14450

13. Cremer, T., & Cremer, C. (2001). Chromosome territories, nuclear architecture and gene regulation in mammalian cells. Nature reviews in genetics, 2, 292–301. doi: 10.1038/35066075

14. Davidson, E. H., Rast, J. P., Oliveri, P., Ransick, A., Calestani, C., Yuh, C. H., … Bolouri, H. (2002). A provisional regulatory gene network for specification of endomesoderm in the sea urchin embryo. Developmental Biology, 246, 162–190. doi: 10.1006/dbio.2002.0635.

15. Davidson, P. L., Guo, H., Wang, L., Berrio, A., Zhang, H., Chang, Y., …Wray, G. A. (2020). Chromosomal-Level Genome Assembly of the Sea Urchin *Lytechinus variegatus* Substantially Improves Functional Genomic Analyses. Genome Biology and Evolution, 12, 1080–1086. doi: 10.1093/gbe/evaa101

16. Dixon, J. R., Selvaraj, S., Yue, F., Kim, A., Li, Y., Shen, Y., … Ren, B. (2012). Topological domains in mammalian genomes identified by analysis of chromatin interactions. Nature, 485, 376–380. doi: 10.1038/nature11082

17. Dowen, J. M., Fan, Z. P., Hnisz, D., Ren, G., Abraham, B. J., Zhang, L. N., … & Young, R. A. (2014). Control of cell identity genes occurs in insulated neighborhoods in mammalian chromosomes. Cell, 159, 374–387. doi: 10.1016/j.cell.2014.09.030

18. Du, Z., Zheng, H., Huang, B., Ma, R., Wu, J., Zhang, X., … Xie, W. (2017). Allelic reprogramming of 3D chromatin architecture during early mammalian development. Nature, 547, 232–235. doi: 10.1038/nature23263.

19. Fedoriw, A. M., Stein, P., Svoboda, P., Schultz, R. M., & Bartolomei, M. S. (2004). Transgenic RNAi reveals essential function for CTCF in H19 gene imprinting. Science, 303, 238–240. doi: 10.1126/science.1090934.

20. Filippova, G. N., Qi, C-F., Ulmer, J. E., Moore, J. M., Ward, M. D., Hu, Y. J., … Lobanenkov, V. V. (2002). Tumor-associated zinc finger mutations in the CTCF transcription factor selectively alter its DNA-binding specificity. Cancer Research, 62, 48–52.

21. Foster, S., Oulhen, N., & Wessel, G. (2020). A single cell RNA sequencing resource for early sea urchin development. Development, 147, dev191528. doi: 10.1242/dev.191528.

22. Franke, M., De la Calle-Mustienes, E., Neto, A., Almuedo-Castillo, M., Irastorza-Azcarate, I., Acemel, R. D., … Gómez-Skarmeta, J. L. (2021). CTCF knockout in zebrafish induces alterations in regulatory landscapes and developmental gene expression. Nature Communications, 12, 5415. doi: 10.1038/s41467-021-25604-5

23. Fudenberg, G., & Nora, E. P. (2021). Embryogenesis without CTCF in flies and vertebrates. Nature Structural & Molecular Biology, 28, 774–776. doi: 10.1038/s41594-021-00662-x

24. Gildor, T., & Ben-Tabou de-Leon, S. (2015). Comparative Study of Regulatory Circuits in Two Sea Urchin Species Reveals Tight Control of Timing and High Conservation of Expression Dynamics. PLoS Genetics, 11, e1005435. doi: 10.1371/journal.pgen.1005435.

25. Heger, P., Marin, B., & Schierenberg, E. (2009). Loss of the insulator protein CTCF during nematode evolution. BMC Molecular Biology, 10, 84. doi: 10.1186/1471-2199-10-84

26. Heger, P., George, R., & Wiehe, T. (2013). Successive gain of insulator proteins in arthropod evolution. Evolution, 67, 2945–2956. doi: 10.1111/evo.12155

27. Heger, P., Marin, B., Bartkuhn, M., Schierenberg, E., & Wiehe, T. (2012). The chromatin insulator CTCF and the emergence of metazoan diversity. Proceedings of the National Academy of Sciences of the United States of America, 109, 17507–17512. doi: 10.1073/pnas.1111941109.

28. Hosoi, S., Sakuma, T., Sakamoto, N., & Yamamoto, T. (2014). Targeted mutagenesis in sea urchin embryos using TALENs. Development, Growth & Differentiation, 56, 92–97. doi: 10.1111/dgd.12099

29. Hug, C. B., Grimaldi, A. G., Kruse, K., & Vaquerizas, J. M. (2017). Chromatin Architecture Emerges during Zygotic Genome Activation Independent of Transcription. Cell, 169, 216–228. doi: 10.1016/j.cell.2017.03.024.

30. Katow, H., Katow, T., Yoshida, H., & Kiyomoto, M. (2021). Involvement of Huntingtin in Development and Ciliary Beating Regulation of Larvae of the Sea Urchin, Hemicentrotus pulcherrimus. International Journal of Molecular Sciences, 22, 5116. doi: 10.3390/ijms22105116

31. Kaushal, A., Mohana, G., Dorier, J., Özdemir, I., Omer, A., Cousin, P., … Gambetta, M. C. (2021). CTCF loss has limited effects on global genome architecture in Drosophila despite critical regulatory functions. Nature Communications, 12, 1011. doi: 10.1038/s41467-021-21366-2

32. Ke, Y., Xu, Y., Chen, X., Feng, S., Liu, Z., Sun, Y., … Liu, J. (2017). 3D Chromatin Structures of Mature Gametes and Structural Reprogramming during Mammalian Embryogenesis. Cell, 170, 367–381. doi: 10.1016/j.cell.2017.06.029.

33. Kim, T. H., Abdullaev, Z. K., Smith, A. D., Ching, K. A., Loukinov, D. I., Green, R. D., … Ren, B. (2007). Analysis of the vertebrate insulator protein CTCF-binding sites in the human genome. Cell, 128, 1231–1245. doi: 10.1016/j.cell.2006.12.048

34. Kim, J., Jimenez, D. S., Ragipani, B., Zhang, B., Street, L. A., Kramer, M., … Ercan, S. (2022). Condensin DC loads and spreads from recruitment sites to create loop-anchored TADs in *C. elegans*. eLife, 11, e68745. doi: 10.7554/eLife.68745

35. Kinjo, S., Kiyomoto, M., Yamamoto, T., Ikeo, K., & Yaguchi, S. (2018). HpBase: A genome database of a sea urchin, *Hemicentrotus pulcherrimus*. Development, Growth & Differentiation, 60, 174–182. doi: 10.1111/dgd.12429

36. Kyrchanova, O., Klimenko, N., Postika, N., Bonchuk, A., Zolotarev, N., Maksimenko, O., & Georgiev, P. (2021). Drosophila architectural protein CTCF is not essential for fly survival and is able to function independently of CP190. Biochimica et Biophysica Acta - Gene Regulatory Mechanisms, 1864, 194733. doi: 10.1016/j.bbagrm.2021.194733

37. Lin, C. Y., & Su, Y. H. (2016). Genome editing in sea urchin embryos by using a CRISPR/Cas9 system. Developmental Biology, 409, 420–428. doi: 10.1016/j.ydbio.2015.11.018

38. Liu, D., Awazu, A., Sakuma, T., Yamamoto, T., & Sakamoto, N. (2019). Establishment of knockout adult sea urchins by using a CRISPR-Cas9 system. Development Growth and Differentiation, 61, 378–388. doi: 10.1111/dgd.12624

39. Lobanenkov, V. V., Nicolas, R. H., Adler, V. V., Paterson, H., Klenova, E. M., Polotskaja, A. V., & Goodwin, G. H. (1990). A novel sequence-specific DNA binding protein which interacts with three regularly spaced direct repeats of the CCCTC-motif in the 5’-flanking sequence of the chicken c-myc gene. Oncogene, 5, 1743–1753.

40. Materna, S. C., Nam, J., & Davidson, E. H. (2010). High accuracy, high-resolution prevalence measurement for the majority of locally expressed regulatory genes in early sea urchin development. Gene Expression Patterns, 10, 177–184. doi: 10.1016/j.gep.2010.04.002.

41. Merkenschlager, M., & Nora, E. P. (2016). CTCF and Cohesin in Genome Folding and Transcriptional Gene Regulation. Annual Review of Genomics and Human Genetics, 17, 17–43. doi: 10.1146/annurev-genom-083115-022339

42. Minokawa, T., Rast, J. P., Arenas-Mena, C., Franco, C. B., & Davidson, E. H. (2004). Expression patterns of four different regulatory genes that function during sea urchin development. Gene Expression Patterns, 4, 449–456. doi: 10.1016/j.modgep.2004.01.009

43. Moore, J. M., Rabaia, N. A., Smith, L. E., Fagerlie, S., Gurley, K., Loukinov, D., … Filippova, G. N. (2012). Loss of maternal CTCF is associated with peri-implantation lethality of *Ctcf* null embryos. PLoS ONE, 7(4). doi: 10.1371/journal.pone.0034915

44. Naumova, N., Imakaev, M., Fudenberg, G., Zhan, Y., Lajoie, B. R., Mirny, L. A., & Dekker, J. (2013). Organization of the mitotic chromosome. Science, 342, 948–53. doi: 10.1126/science.1236083

45. Ochiai, H., Fujita, K., Suzuki, K. I., Nishikawa, M., Shibata, T., Sakamoto, N., & Yamamoto, T. (2010). Targeted mutagenesis in the sea urchin embryo using zinc-finger nucleases. Genes to Cells, 15, 875–885. doi: 10.1111/j.1365-2443.2010.01425.x

46. Ohlsson, R., Renkawitz, R., & Lobanenkov, V. (2001). CTCF is a uniquely versatile transcription regulator linked to epigenetics and disease. Trends in Genetics, 17, 520–527. doi: 10.1016/s0168-9525(01)02366-6

47. Oliveri, P., & Davidson, E. H. (2004). Gene regulatory network controlling embryonic specification in the sea urchin. Current Opinion in Genetics & Development, 14, 351–360. doi: 10.1016/j.gde.2004.06.004.

48. Oliveri, P., Tu, Q., & Davidson, E. H. (2008). Global regulatory logic for specification of an embryonic cell lineage. Proceedings of the National Academy of Sciences of the United States of America, 105, 5955–5962. doi: 10.1073/pnas.0711220105.

49. Oomen, M. E., Hansen, A. S., Liu, Y., Darzacq, X., & Dekker, J. (2019). CTCF sites display cell cycle-dependent dynamics in factor binding and nucleosome positioning. Genome Research, 29, 236–249. doi: 10.1101/gr.241547.118

50. Palozola, K. C., Donahue, G., Liu, H., Grant, G. R., Becker, J. S., Cote, A., … Zaret, K. S. (2017). Mitotic transcription and waves of gene reactivation during mitotic exit. Science, 358, 119–122. doi: 10.1126/science.aal4671

51. Parelho, V., Hadjur, S., Spivakov, M., Leleu, M., Sauer, S., Gregson, H. C., … Merkenschlager, M. (2008). Cohesins Functionally Associate with CTCF on Mammalian Chromosome Arms. Cell, 132(3), 422–433. doi: 10.1016/j.cell.2008.01.011

52. Racko, D., Benedetti, F., Dorier, J., & Stasiak, A. (2019). Are TADs supercoiled?. Nucleic Acids Research, 47, 521–532. doi: 10.1093/nar/gky1091

53. Rast, J. P. (2000) Transgenic manipulation of the sea urchin embryo. Methods in Molecular Biology, 136, 365–373. doi: 10.1385/1-59259-065-9:365

54. Satou, Y., Nakamura, R., Yu, D., Yoshida, R., Hamada, M., Fujie, M., … Satoh, N. (2019). A Nearly Complete Genome of *Ciona intestinalis* Type A (*C. robusta*) Reveals the Contribution of Inversion to Chromosomal Evolution in the Genus *Ciona*. Genome Biology and Evolution, 11, 3144–3157. doi: 10.1093/gbe/evz228

55. Sea Urchin Genome Sequencing Consortium (2006). The genome of the sea urchin *Strongylocentrotus purpuratus*. Science, 314, 941–952. doi: 10.1126/science.1133609.

56. Sexton, T., Schober, H., Fraser, P., & Gasser, S. M. (2007). Gene regulation through nuclear organization. Nature structural & molecular biology, 14, 1049–1055. doi: 10.1038/nsmb1324

57. Szabo, Q., Bantignies, F., & Cavalli, G. (2019). Principles of genome folding into topologically associating domains. Science advances, 5, eaaw1668. doi: 10.1126/sciadv.aaw1668

58. Van Bortle, K., Nichols, M. H., Li, L., Ong, C. T., Takenaka, N., Qin, Z. S., & Corces, V. G. (2014). Insulator function and topological domain border strength scale with architectural protein occupancy. Genome Biology, 15, R82. doi: 10.1186/gb-2014-15-5-r82

59. Wendt, K. S., Yoshida, K., Itoh, T., Bando, M., Koch, B., Schirghuber, E., … Peters, J. M. (2008). Cohesin mediates transcriptional insulation by CCCTC-binding factor. Nature, 451(7180), 796–801. doi: 10.1038/nature06634

60. Wike, C. L., Guo, Y., Tan, M., Nakamura, R., Shaw, D. K., Díaz, N., … Cairns, B. R. (2021). Chromatin architecture transitions from zebrafish sperm through early embryogenesis. Genome Research, 31, 981–994. doi: 10.1101/gr.269860.120.

61. Yajima, M., Fairbrother, W. G., & Wessel, G. M. (2012). ISWI contributes to ArsI insulator function in development of the sea urchin. Development, 139, 3613–3622. doi: 10.1242/dev.081828

62. Yan, Y., Cummings, C. A., Sutton, D., Yu, L., Castro, L., Moore, A. B., … Dixon, D. (2016). Immunogold electron microscopy and confocal analyses reveal distinctive patterns of histone H3 phosphorylation during mitosis in MCF-7 cells. Genes Chromosomes & Cancer, 55, 397–406. doi: 10.1002/gcc.22343.

