## Supplementary material for "The Crucial Role of CTCF in Mitotic Progression during Early Development of Sea Urchin": Table S1

### Supporting information

(a)

| **Stage** | **GO Term(Positive Correlation with *SpCTCF*)** |
|---|---|
| Early brastula | Establishment or maintenance of cell polarity<br>Regulation of phosphate metabolic process<br>Regulation of protein modification process<br>mRNA processing |
| Hatched brastula | Establishment of mitotic spindle localization<br>Establishment or maintenance of cell polarity<br>mRNA transport<br>Translation<br>Negative regulation of nucleobase-containing compound<br>Metabolic process<br>Chromatin remodeling<br>DNA conformation change<br>Regulation of cell cycle<br>mRNA splicing, via spliceosome<br>Ribonucleoprotein complex biogenesis<br>Protein-containing complex organization<br>Regulation of dna-templated transcription<br>G protein-coupled receptor signaling pathway |
| Mesenchyme brastula | Establishment of mitotic spindle localization<br>Regulation of protein kinase activity<br>Nucleosome assembly<br>Regulation of cell cycle process<br>Translation<br>Electron transport chain<br>mRNA processing<br>DNA conformation change<br>RNA splicing<br>Protein folding<br>Regulation of transcription by RNA polymerase II |
| Early gastrulation | DNA replication-dependent chromatin assembly<br>Positive regulation of cell cycle phase transition<br>Negative regulation of DNA metabolic process<br>Chromosome condensation<br>Translational elongation<br>One-carbon metabolic process<br>Histone lysine methylation<br>Nucleosome assembly<br>Regulation of protein kinase activity<br>Regulation of mitotic cell cycle phase transition<br>Protein acetylation<br>Nucleoside monophosphate biosynthetic process<br>Cell division<br>Cell cycle<br>Regulation of cellular component organization<br>mRNA processing<br>Ribonucleoprotein complex biogenesis<br>RNA splicing<br>Organelle assembly<br>Cellular response to DNA damage stimulus<br>Regulation of dna-templated transcription |
| Late gastrulation | Regulation of alternative mrna splicing, via spliceosome<br>Chromosome condensation<br>CTP biosynthetic process<br>Translational elongation<br>Histone lysine methylation<br>Regulation of protein kinase activity<br>Nucleosome assembly<br>Cytoplasmic translation<br>Regulation of mitotic cell cycle<br>Histone acetylation<br>Regulation of cell cycle process<br>Translational initiation<br>Cell division<br>Cell cycle<br>mRNA processing<br>Ribosome biogenesis<br>Negative regulation of macromolecule metabolic process<br>Regulation of transcription by RNA polymerase II |

(b)

| Stage | GO Term(Negative Correlation with *SpCTCF*) |
|---|---|
| Early brastula | - |
| Hatched brastula | - |
| Mesenchyme brastula | - |
| Early gastrulation | Ion transmembrane transport |
| Late gastrulation | - |

(c)

| Stage | GO Term(Non Correlation with *SpCTCF*) |
|---|---|
| Early brastula | - |
| Hatched brastula | - |
| Mesenchyme brastula | - |
| Early gastrulation | - |
| Late gastrulation | - |

**Table S1**. (a)Biological process of transcript which have positive correlation with *SpCTCF* Enriched upstream functions as transcript functions positively correlated with *SpCTCF*. Red text indicates cell cycle-related functions. Number of transcript in each stage : (EB = 284, HB = 510, MB = 474, EG = 504, LG = 505) (b) Biological process of transcript which have negative correlation with *SpCTCF* Enriched upstream functions as transcript functions negatively correlated with *SpCTCF*. Number of transcript : (EB, HB, MB, EG, LG = 99, 105, 146, 173, 223) (c) Biological process of transcript which have no correlation with *SpCTCF.* There were no specifically enriched functions in the transcripts that did not correlate with *SpCTCF*. Number of transcript : (EB, HB, MB, EG, LG = 284, 510, 474, 504, 505) Single cell data were obtained from the GEO database (GSE149221) and analyzed using EB to LG stage data. Data filtering was performed according to the original paper from which the data were obtained (Foster et al., 2020). Correlations between each transcript and SpCTCF expression levels were calculated to extract transcripts correlated with SpCTCF and a test of the mother correlation coefficient was performed (p = 0.00001). In addition to groups with positive correlations groups with negative correlations and groups with small absolute values of correlation coefficients were extracted for comparison. The number of groups with small absolute values of correlation coefficients was kept equal to the number of groups with positive correlations at eachzdevelopmental stage. Enrichment analysis was performed using Gene ontology resource(http://geneontology.org) and focused on the top biological process function enriched in each extracted group.
